# Modeling spheroid assembly dynamics and mechanical stress generation in magnetic-based biofabrication

**DOI:** 10.64898/2026.06.05.730349

**Authors:** Maya Guilliams, Konstantinos Ioannidis, Karolina Z. Dąbrowska, Marta Tosini, Dimitrios Lefas, Gianpaolo Serino, Dimitrios Sakellariou, Ioannis Papantoniou, Bart Smeets

## Abstract

Magnetic biofabrication enables rapid assembly of multicellular spheroids but still lacks a basis for predictive control over structure and mechanical environment. Here, we combine experiments and an individual spheroid-based model to study magnetic assembly of periosteum-derived spheroids. Spheroids are treated as discrete particles interacting through magnetic forces, contact mechanics, and interfacial friction, with parameters obtained from independent measurements. This model quantitatively captures assembly dynamics arising from magnetic force patterns and viscous drag with the well surface. The spatial distribution of magnetic forces, determined by magnet geometry and positioning, predicts the size and morphology of magnetic assembloids, including disk- and ring-like structures. Magnetic assembly further generates heterogeneous compressive stresses that depend on magnet geometry and spheroid number. Radial stresses arise collectively through inter-spheroid interactions, whereas vertical stresses are mainly determined by magnetic loading of individual spheroids. These results establish a minimal physical framework for magnetic biofabrication and provide a basis for predictive control of both tissue structure and mechanical microenvironment.

## 1. Introduction

Biofabrication technologies provide the means to spatially control deposition of cells, spheroids, microtissues and biomaterials, into precise and “as designed” configurations, for in vitro modeling and regenerative medicine applications [1]. These technologies can be broadly distinguished into bioprinting and bioassembly processes [2]. Bioassembly processes can be viewed as bottom up bio-fabrication technologies where living modules such as multicellular spheroids, microtissues, organoids and micromaterials serve as building blocks for engineering larger assembloid tissues [3]. In bottom-up biofabrication strategies, scaffold-free methodologies are explored, where only cells and living modules are used to create 3D structures in the absence of supporting materials, bypassing concerns like scaffold choice, degradation rate and toxicity of degradation products, immunogenicity, host inflammatory responses and mechanical mismatch with the native tissue [4]. In this context, magnetic-based biofabrication has gained attention due to its ability to enable contactless, bottom-up, scaffold-free biofabrication techniques [5]. Magnetic nanoparticles (MNPs) have been extensively used in biomedical applications because of their biocompatibility and remote actuation in magnetic fields [4]. In this approach, cells or spheroids are incubated with MNPs, allowing external magnetic fields to guide their assembly into prescribed configurations while also enabling remote mechanical stimulation [6, 7, 8, 9]. Superparamagnetic nanoparticles are predominantly used, enabling reversible ON/OFF actuation states that can influence the patterning, 3D shape, and mechanical stimulation of the resulting constructs [10, 11]. While this method enables rapid spheroid assembly and organization, control over the final structure remains limited, as the process is largely passive and governed by the interplay between magnetic forces and physical interactions at the well surface.

Computational models for magnetic biofabrication should capture the essential physics of spheroid assembly while remaining tractable for large spheroid populations. Existing biofabrication models often operate at the cell scale, including Cellular Potts, kinetic Monte Carlo, and center-based approaches [12, 13, 14]. Although these methods can represent cell behavior and cell-cell interactions in detail, they become computationally expensive for systems containing hundreds to thousands of spheroids. At the other extreme, continuum and phase-field models describe tissue-scale behavior but are less suited to resolving discrete spheroid interactions during early assembly and often rely on parameters that are difficult to measure experimentally [15, 16]. There is therefore a need for a spheroid-scale framework that directly links magnetic forces, interfacial mechanics, and assembly dynamics. Magnetic assembly also generates mechanical stresses within the developing construct through both direct magnetic loading and inter-spheroid interactions. These stresses can influence cell behavior, differentiation, and tissue maturation [17, 18, 19, 20]. However, current models of magnetic biofabrication rarely quantify how stresses emerge during assembly or how they depend on experimental conditions. This requires identifying the minimal set of physical mechanisms, including magnetic forces, contact mechanics, and friction, that govern both spheroid organization and the resulting mechanical environment.

In this work, we introduce an individual spheroid-based model in which spheroids are treated as discrete particles interacting through magnetic forces, contact mechanics, and well surface friction. The model is parameterized using mainly experimentally accessible quantities and is applied to simulate the magnetic assembly of periosteum-derived cell spheroids. We show that this minimal framework captures the observed assembly dynamics, spatial organization, and emergence of collective stress patterns. In addition, we use the model to quantify how magnet geometry, spheroid number, and magnetic loading determine internal stress distributions, providing a basis for the predictive control of both structure and mechanical conditions in magnetic biofabrication.

## 2. Results

### Magnetic assembly across controlled spheroid conditions

We seeded human periosteum-derived cells (hPDCs) as single cells in non-adherent well-plates to form multicellular spheroids. After formation, these spheroids were incubated with magnetic nanoparticles (MNPs), resulting in magnetized spheroids. These spheroids were then transferred to a magnetic plate with a permanent magnet positioned underneath each well (Fig. 1A). Immediately after, the aggregation process of spheroids assembling on top of the magnet was imaged (Fig. 1C), with the resulting aggregates of spheroids hereafter referred to as “assembloids”. Spheroids cultured for 3 and 7 days in chondrogenic differentiation medium were included to compare two early stages of maturation. Previous characterization of this differentiation system showed that major biological changes already occur during the first week of culture, with progression from a compact cellular spheroid toward a more differentiated microtissue state, accompanied by increasing cartilage-like extracellular matrix (ECM) deposition [21]. These two conditions therefore represent distinct states, transitioning from a more cell-dominated assembloid at day 3 to a more ECM-rich, tissue-like spheroid at day 7 (Fig. 1B). Still, the day 3 and day 7 spheroids had comparable mechanical stiffness as measured by nanoindentation, with effective elastic moduli of 361 ± 240 Pa and 254 ± 101 Pa, respectively (Fig. 1D-E). In addition, we varied spheroid size using two well formats. A400 wells produce smaller spheroids with a radius of approximately 100 µm, whereas A800 wells produce larger spheroids with a radius of approximately 175 µm (Fig. 1B,F). For both conditions, the same total number of cells and total amount of MNPs are used, resulting in either many small spheroids (A400) or fewer large spheroids (A800). Consequently, the available MNPs are distributed over fewer spheroids in the A800 condition, such that each spheroid experiences a larger total magnetic force. These conditions therefore provide control over spheroid size, mechanical state, and magnetic forcing, while keeping all other system parameters fixed.

**Figure 1:**
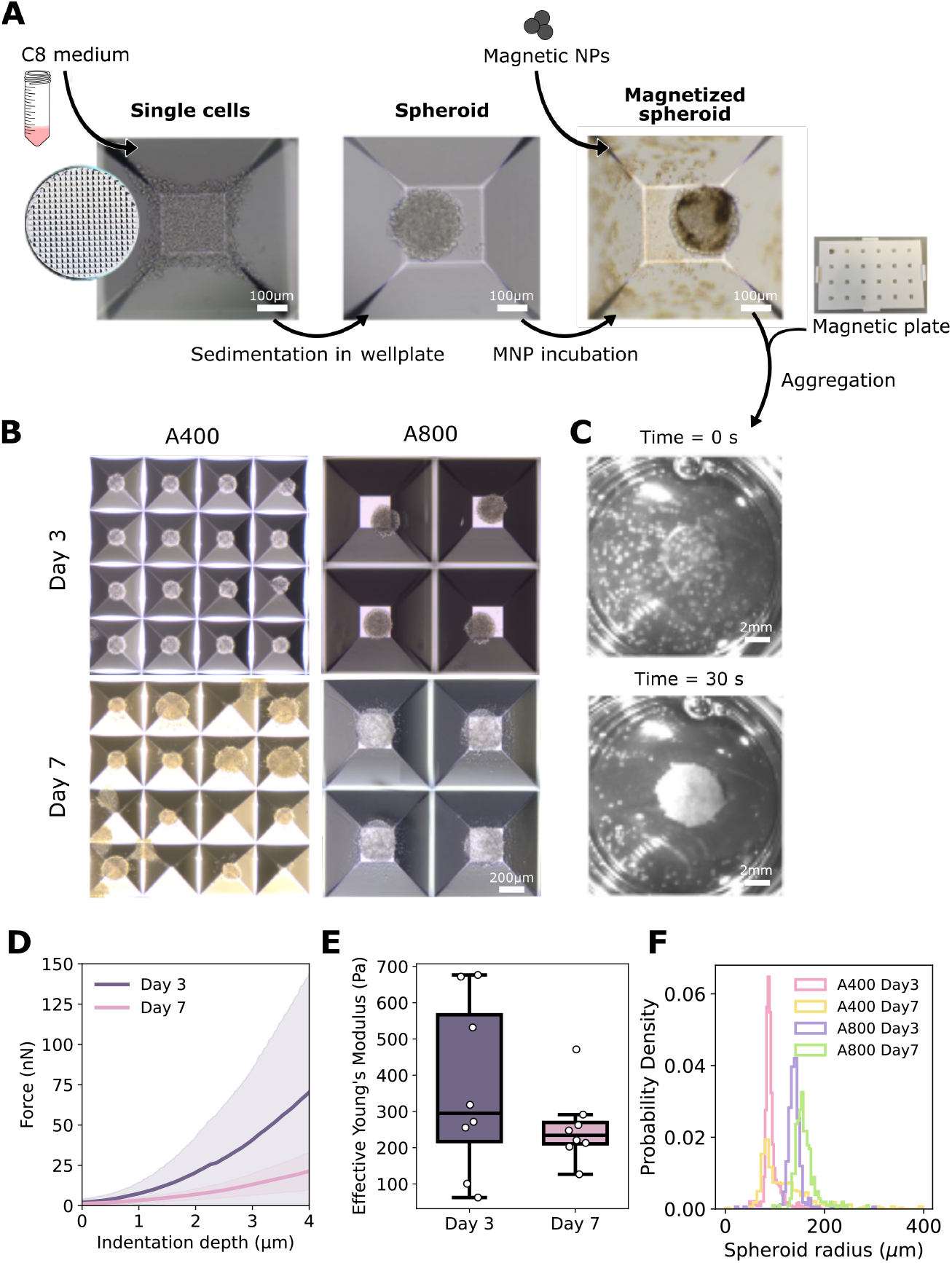
**(A)** Schematic of the experimental setup. Cells sediment in microwells and self-assemble into multicellular spheroids. Magnetic nanoparticles (MNPs) are added one day prior to assembly, after which magnetized spheroids are transferred to non-adherent wells and exposed to a magnetic field to induce assembly. **(B)** Microscopy images of spheroids generated in different well formats (A400 and A800) at two maturation stages (Day 3 and Day 7). **(C)** Microscopy images immediately after magnetic field application and after 30 s, illustrating magnetic assembly into an assembloid. **(D)** Average force–indentation curves (approach phase) for A800 spheroids cultured in C8 medium for 3 or 7 days, aligned at the contact point. Individual curves were averaged and smoothed using a Savitzky– Golay filter. Solid lines represent the mean force as a function of indentation depth, and shaded regions represent ± standard deviation. Data were obtained from 8 spheroids per time point, comprising 60 indentations at day 3 and 70 indentations at day 7. **(E)** Effective Young’s modulus of spheroids at Day 3 and 7, determined by nanoindentation. Each data point represents a single spheroid (n=8 per time point), with values of effective elastic modulus averaged across all indentations performed on that spheroid. **(F)** Spheroid size and number for the A400 (Day 3: n=1159; Day 7: n=409) and A800 (Day 3: n=294; Day 7: n=261) conditions.

### Individual-based model of magnetically driven spheroids

Based on this experimental setup, we developed an individual-based computational model (IbM), formulated as an overdamped Discrete Element Method (DEM), in which each spheroid is represented as a single spherical particle. The motion of each spheroid is governed by gravity, Hertzian repulsive interactions with other spheroids and the well surface, viscous drag with the well surface, and magnetic forces arising from incorporated nanoparticles (Fig. 2A). To compute these forces, we resolve the spatial magnetic field and couple it to a magnetization curve describing the field-dependent response of the nanoparticles (Fig. 2B-C). The resulting forces are combined into an overdamped equation of motion, which is integrated to track spheroid trajectories during assembly. Our simulations mimic the experimental procedure described above, with spheroids initialized above the magnet and subsequently sedimenting and aggregating under the combined influence of gravity and magnetic forces.

**Figure 2:**
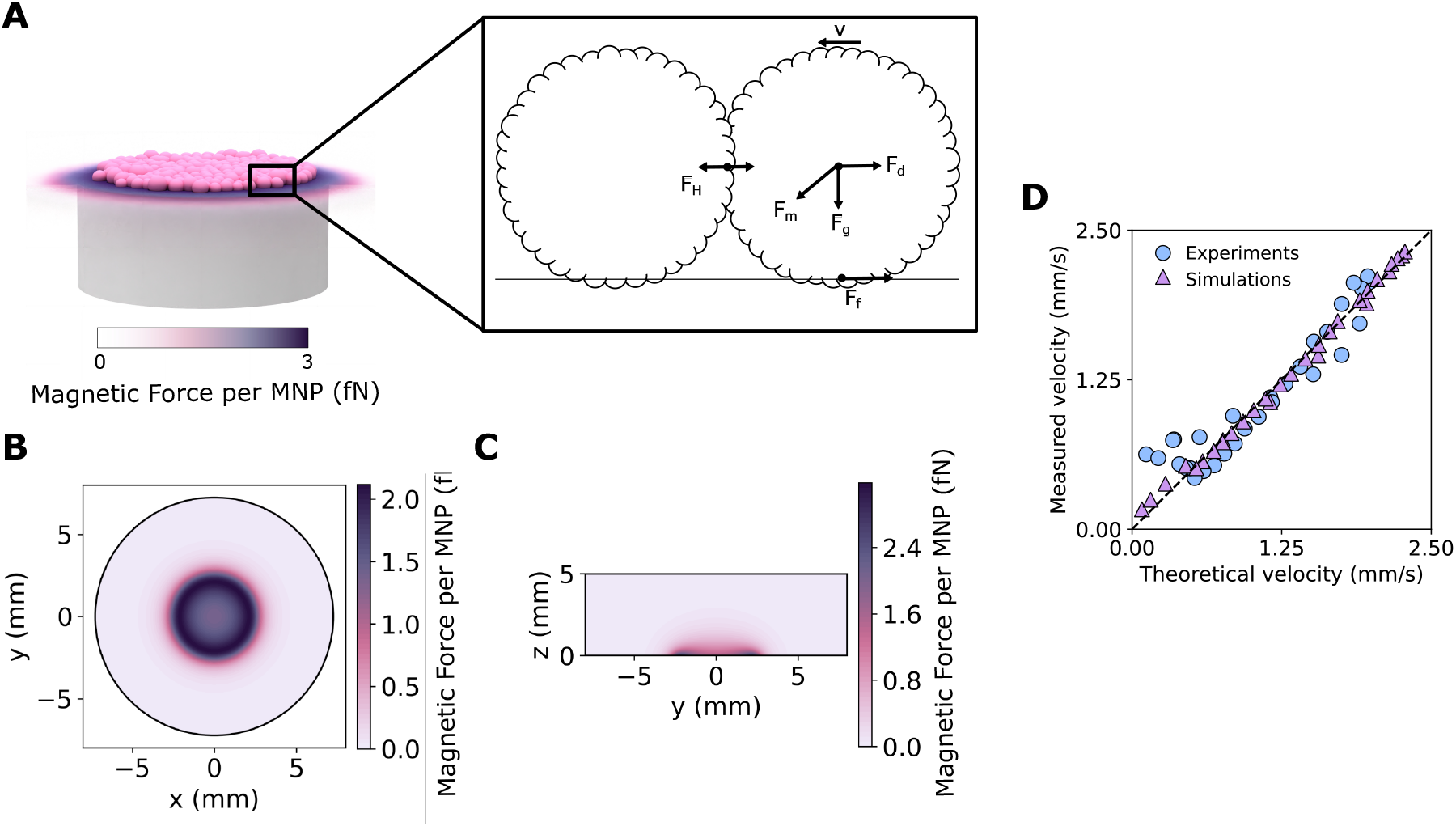
**(A)** Individual spheroid-based model of magnetically guided spheroid motion. The equations of motion include the magnetic force **F**_*m*_, gravitational force **F**_*g*_, drag force **F**_*d*_, friction force with the well surface **F**_*f*_, and contact force between colliding spheroids **F**_*H*_, with spheroid velocity **v. (B)** Top view of the magnetic force magnitude acting on an MNP at the bottom plane of the well, showing the radial force distribution in the x–y plane. **(C)** Side-view heat map of the magnetic force magnitude acting on an MNP in the central vertical (x–z) plane of the well. **(D)** Comparison of radial spheroid velocities from experiments (834 spheroids tracked individually) and simulations (velocity during the first second for 10615 spheroids), together with the theoretical prediction *v*_*t*_ = *F*_*m,t*_*/*(6*πRη* +*c*_*a*_*S*) using a fitted value of *c*_*a*_ = 33.24 Pa s/mm, for spheroids located 1.5–6 mm from the center; data are binned over 25 bins.

The experimental system provides direct access to the key physical inputs governing assembly. Spheroid size and mechanical stiffness are measured independently, and magnetic forces can be estimated from the spatial magnetic field and a prescribed magnetization curve, assuming a fixed fraction of incorporated MNPs in spheroid material [22]. This leaves interfacial interactions between spheroids and the well surface as the primary unknown. We model these interactions as effective wet interfacial friction with a coefficient *c*_*a*_, acting over the contact area. To estimate *c*_*a*_ from experiments, we compare the measured tangential spheroid velocity with the theoretical velocity obtained from a balance between tangential magnetic forcing *F*_*m,t*_ and viscous resistance (Appendix A). The latter consists of the Stokes drag and an additional interfacial contribution proportional to the contact area *S*, yielding an effective resistance 6*πRη* + *c*_*a*_*S*, with spheroid radius *R* and medium viscosity *η*. The contact area is estimated from Hertzian contact theory as

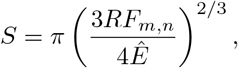

with normal magnetic force *F*_*m,n*_ and effective elastic modulus *Ê*. The parameter *c*_*a*_ is then obtained as the single free fit parameter by matching the theoretical and experimentally measured velocities across all simulated conditions, yielding *c*_*a*_ ≈ 33 Pa s*/*mm, Fig. 2D.

### Model reproduces assembly dynamics and structure

Magnetic assembly occurs rapidly in both experiments and simulations (Fig. 3A). Spheroids are collected at the magnet surface within seconds, with approximately 50% of the population reaching the magnet within 0.5 s and complete assembly occurring on the order of 30 s. The IbM simulations reproduce these dynamics and capture the experimentally observed aggregation pathways, with close visual agreement between experimental and simulated configurations (Fig. 3A and Supplementary Fig. S1-S2). We quantified assembloid size and assembly dynamics across eight conditions (Fig. 3B-J). At fixed total magnetic nanoparticle (MNP) content and conserved total cell volume, increasing spheroid size leads to more compact assemblies, reflected by a reduced final assembloid area (Fig. 3B). Day 7 spheroids form larger assembloids; however, this increase coincides with an increase in individual spheroid size, consistent with increased spheroid volume due to ECM production [22]. In contrast, increasing MNP loading from 30 to 60 µg has only a moderate effect on the final assembloid area, indicating that assembly geometry is relatively insensitive to magnetic forcing within this range (Fig. 3B,C). Because the precise number of spheroids participating in assembloid formation varies between samples and strongly affects the final assembloid area, we compared experiments and simulations using the relative area change between the 30 and 60 µg conditions (Fig. 3C). Both experiments and simulations show only limited additional compaction at higher MNP loading, suggesting that stronger magnetic forces primarily increase compressive stresses within the assembloid rather than causing substantial spheroid rearrangements.

**Figure 3:**
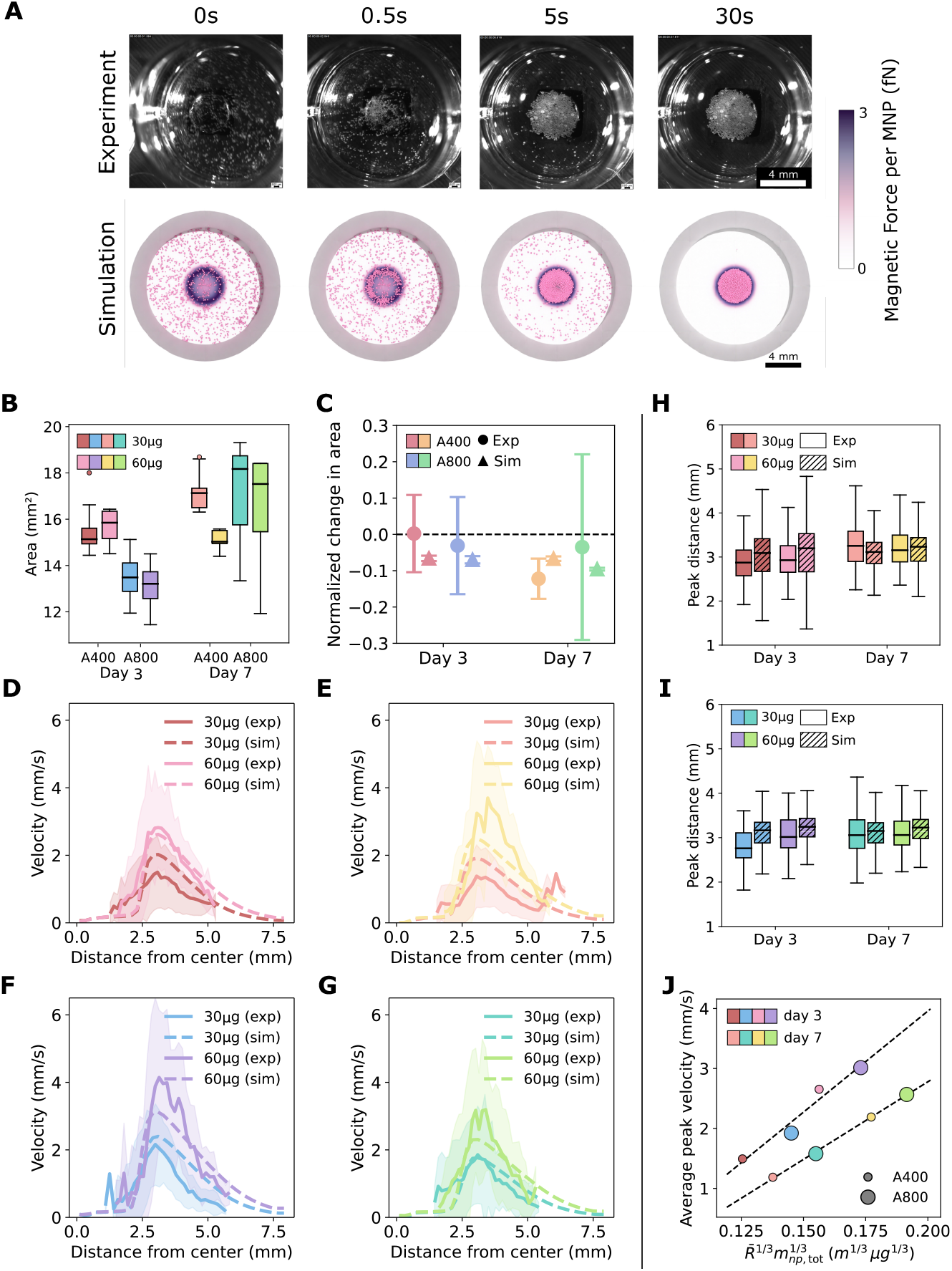
**(A)** Time-lapse comparison of magnetic spheroid assembly in experiments and simulations during the first 30 s of aggregation. The color scale in the simulations indicates the magnetic force per MNP. **(B)** Final assembloid area (convex hull area of the largest spheroid cluster) for varying spheroid size (A400 and A800), MNP loading (30 µg and 60 µg), and spheroid maturation state (Day 3 and Day 7). **(C)** Normalized change in final assembloid area of Day 3 spheroids between the 30 µg and 60 µg MNP conditions, comparing experiments and simulations. **(D–G)** Radial velocity profiles of tracked spheroids as a function of distance from the magnet center for small spheroids at Day 3 **(D)**, large spheroids at Day 3 **(E)**, small spheroids at Day 7 **(F)**, and large spheroids at Day 7 **(G). (H–I)** Position of the peak velocity relative to the magnet center for A400 spheroids at Day 3 **(H)** and Day 7 **(I). (J)** Scaling of the peak velocity, averaged over distances between 2.5 and 3.5 mm from the magnet center (see H,I), with the theoretically predicted scaling at high MNP loading, 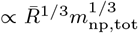 (see Appendix B), where 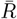 is the average spheroid radius for each condition and *m*_np,tot_ is MNP loading. Dashed lines indicate linear fits for Day 3 and Day 7. Number of independent assembloids (n) and total tracked spheroids (N): A400 Day 3-30 µg (n=5, N=130), 60 µg (n=4, N=104); Day 7-30 µg (n=5, N=130), 60 µg (n=5, N=117). A800 Day 3-30 µg (n=4, N=105), 60 µg (n=3, N=78); Day 7-30 µg (n=3, N=79), 60 µg (n=4, N=104).

To investigate aggregation dynamics, we tracked the radial velocity of individual spheroids (Fig. 3 D–G). Velocities increase as spheroids approach the magnet, reaching a maximum at approximately 3 mm from the center, coinciding with the peak in tangential magnetic force (Fig. 3H–I). The radial velocity increases with MNP loading and shows a weak increase with spheroid size. In the IbM, this observation is consistent with the viscous surface drag model introduced above, for which the interfacial traction was independently estimated from experimental measurements. Combining this with Hertzian contact mechanics, where indentation is set by the normal magnetic force, yields a scaling of the radial velocity (tangential to the well surface) 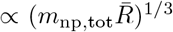 (Appendix B), with nanoparticle load *m*_np,tot_ and typical spheroid radius 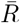, in agreement with the observed scaling of the average peak velocity (Fig. 3J). Close to the magnet center, spheroid velocities sharply decrease to near-zero values, as the in-plane magnetic forces drop in magnitude away from the edge, and due to densification and steric interactions with already assembled spheroids. The consequences of this effect are examined further in the subsequent section. The IbM captures both the spatial organization of the final assembloids and their assembly dynamics, indicating that the system behaves as a driven granular assembly in a centrally directed force field, with dynamics controlled mainly by viscous spheroid-surface friction.

### Ring-like assemblies result from reversal of tangential magnetic forces

At strong magnetic fields and sufficiently small separations between the magnet surface and the well surface, spheroids assemble into stable ring-like structures characterized by depletion near the magnet center and accumulation at a finite radial distance (Fig. 4C). This behavior originates directly from the spatial structure of the magnetic force field. Calculating the tangential magnetic force across the well surface reveals that, at small magnet–well surface separations, the force above the central region of the magnet reverses direction and points outward toward the magnet edge (Fig. 4A). This outward-directed region is surrounded by the inward-directed magnetic attraction outside the magnet perimeter, creating a radial trapping zone that stabilizes spheroids near the magnet edge. As the magnet–well surface separation increases, this outward-directed region progressively disappears, suppressing ring formation.

**Figure 4:**
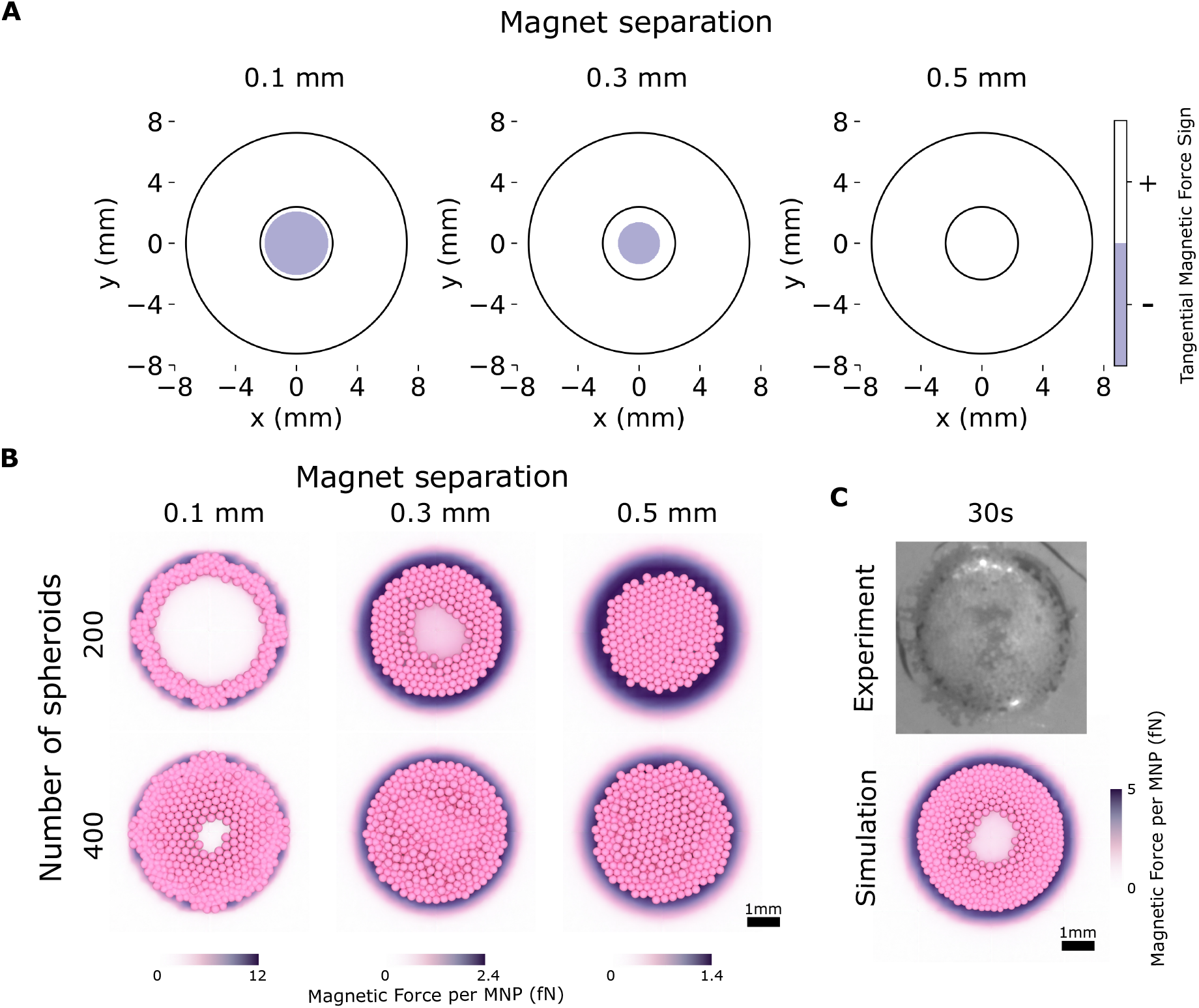
**(A)** Sign of the inward radial component of the magnetic force acting on spheroids for different distances between the magnet and well surfaces (0.1 mm, 0.3 mm, and 0.5 mm). Positive values indicate forces directed toward the magnet center, whereas negative values indicate forces directed toward the magnet edge. **(B)** Simulated endpoint assembloid morphologies for different spheroid numbers (*N*_*s*_ = 200 and *N*_*s*_ = 400). The background color scale indicates the magnetic force magnitude per MNP. **(C)** Comparison of experimentally observed and simulated assembloid structures after 30 s for a magnet–well surface separation of 0.25 mm.

IbM simulations incorporating this magnetic force landscape reproduce the appearance of ring-like assembloids (Fig. 4B). However, the outward tangential force above the magnet is substantially weaker than the inward-directed force experienced by spheroids outside the magnet boundary. Hence, as spheroid number increases, the collective inward pushing by outer spheroids destabilizes the ring and eventually drives collapse toward the magnet center. The simulated assembloids qualitatively match the experimentally observed ring structures for a magnet–well surface separation of 0.25 mm (Fig. 4C), demonstrating that geometry-dependent reversal of the tangential magnetic force is sufficient to explain the formation and collapse of ring-like aggregation patterns.

### Magnet geometry determines spatial stress distributions in spheroid assemblies

We next use the IbM to predict mechanical stress distribution in spheroid assembloids formed under different magnet geometries. Spheroids with identical total MNP content are seeded above cylindrical, ring-shaped, and square magnets and allowed to assemble under magnetic forcing (Fig. 5A–C and Supplementary Fig. S3). For each configuration, we compute a coarse-grained inter-spheroid stress tensor using the virial stress theorem and extract the radial and vertical components for each spheroid. Spatial maps of the radial stress, averaged over 50 independent simulations, are shown in Fig. 5A’–C’. In all cases, stresses are negative, indicating compression, and increase in magnitude with increasing MNP loading (Fig. 5A”–C”). For the cylindrical magnet (Fig. 5A-A”), the radial stress plateaus on the surface of the magnet. The ring-shaped magnet (Fig. 5B-B”) produces a maximum in radial stress near the outer edge. For the square magnet (Fig. 5C-C”), the radial stress is comparatively uniform across the assembly. These results show that magnet geometry directly controls both the magnitude and spatial distribution of compressive stresses within spheroid assemblies, with cylindrical magnets producing the highest central stresses.

**Figure 5:**
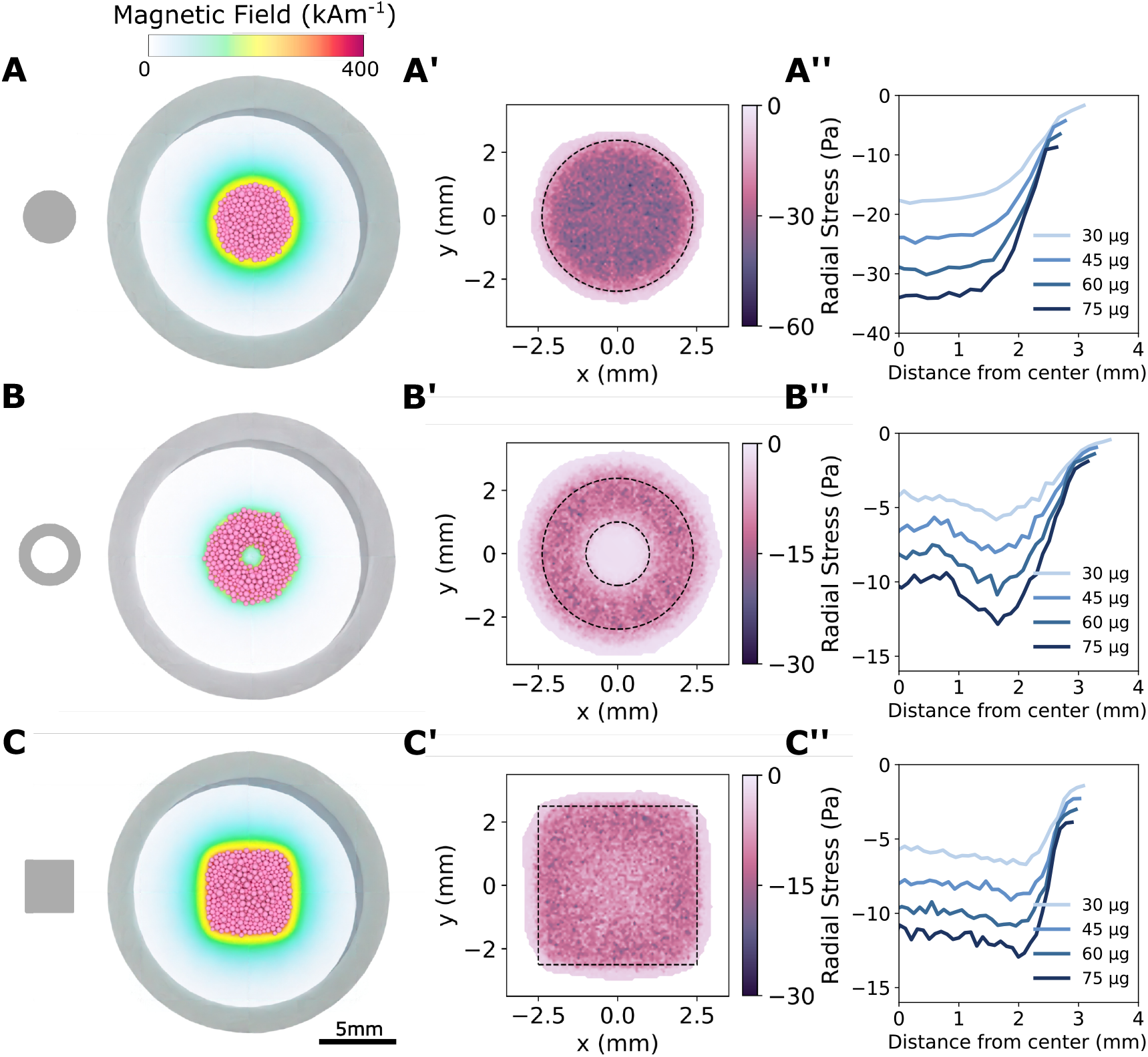
**(A–C)** Magnetic field strength generated by cylindrical **(A)**, ring-shaped **(B)**, and square **(C)** magnets used to drive spheroid assembly. **(A’–C’)** Spatial distributions of the averaged (virial) radial stress within the assembled spheroid assembloids. **(A”–C”)** Averaged radial stress profiles as a function of distance from the center for increasing MNP content (30–75 µg). All quantities are averaged over 50 independent repeats. All simulations were performed with *N*_*s*_ = 400, 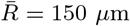, and *Ê*= 361 Pa. The cuboidal magnet had a side length of 4.76 mm, while the ring magnet had an inner radius of 1.0 mm, outer radius of 2.35 mm, and length of 4.76 mm.

### Spheroid number and magnetic loading determine collective stress pattern

We further examine how spheroid number and MNP loading determine stress distributions in assemblies formed on cylindrical magnets (Fig. 6 and Supplementary Fig. S4). For each condition, the total MNP content is conserved, such that increasing the number of spheroids reduces the MNP content per spheroid. Radial stress maps (Fig. 6A) show that compressive stress initially increases with spheroid number, as inter-spheroid interactions generate larger inward forces and increase central stress. This increase saturates between 400 and 800 spheroids, where additional spheroids located near the periphery contribute little tangential force and the available MNPs are distributed over more spheroids. Consistent with this, the mean radial stress peaks at intermediate spheroid numbers (Fig. 6B), reflecting a balance between collective force transmission and reduced magnetic loading per spheroid. In contrast, the vertical (normal) stress is primarily set by the magnetic force acting on individual spheroids and therefore decreases with increasing spheroid number, with the highest values observed at low spheroid counts (Fig. 6E). The relative contribution of radial and vertical stress thus depends on spheroid number. At the single-spheroid level, radial stress exhibits a bimodal distribution, with weakly stressed spheroids at the periphery and strongly compressed spheroids near the center (Fig. 6C), whereas vertical stress remains unimodal (Fig. 6F). The radial stress distribution is most uniform at intermediate spheroid numbers (400–800), where the magnet surface is approximately fully covered (Fig. 6D,G).

**Figure 6:**
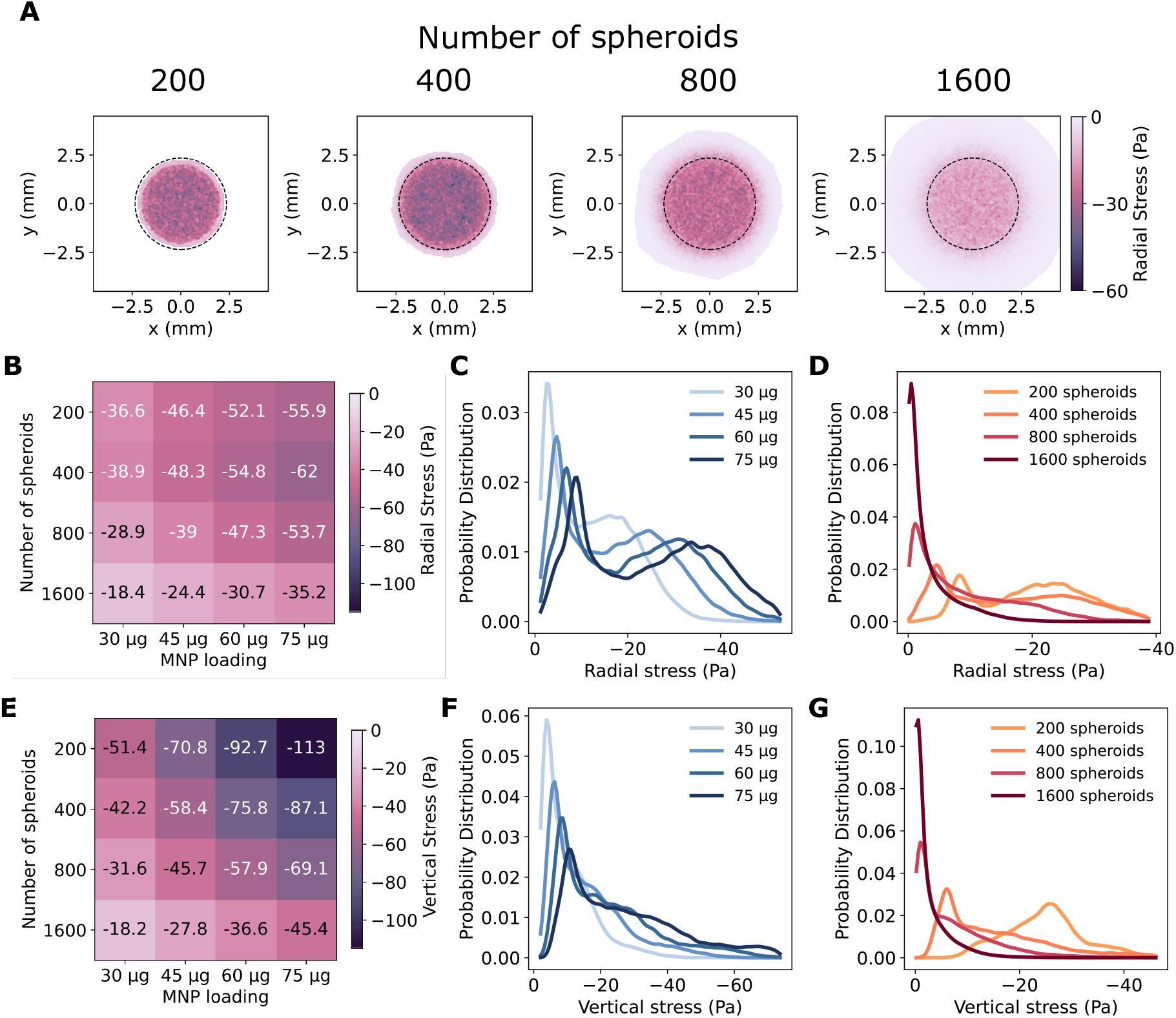
**(A–A” ‘)** Heatmap of radial stress for assemblies composed of 200, 400, 800, and 1600 spheroids, respectively, with the total MNP content held constant. **(B)** Mean radial stress as a function of spheroid number and MNP loading. **(C)** Probability distribution of radial stress per spheroid for different MNP loadings. **(D)** Radial stress distribution for varying spheroid numbers. **(E)** Mean vertical (normal) stress as a function of spheroid number and MNP loading. **(F)** Probability distribution of vertical stress per spheroid for different MNP loadings. **(G)** Vertical stress distribution for varying spheroid numbers. All quantities are averaged over 50 independent repeats.

## 3. Discussion

Our results show that assembly structure and dynamics are governed by a balance between magnetic driving and interfacial frictional interactions at the well surface. Across all conditions, assembly occurs rapidly and reproducibly and can be quantitatively represented by an overdamped individual-based model with minimal assumptions. The observed agreement between experiments and simulations indicates that the system can be understood as a driven particulate assembly, in which spheroids behave as discrete, interacting units whose motion is controlled by effective physical interactions rather than detailed biological complexity, in contrast to later integration and fusion stages that involve additional biological processes.

The assembly dynamics are well described by an effective interfacial drag framework, in which spheroid motion results from a balance between magnetic forces and well surface-mediated dissipation. The scaling of radial velocity with spheroid size and magnetic loading is consistent with a picture in which wet sliding resistance is proportional to the contact area, coupled to Hertzian contact mechanics that set the normal load. This leads to a simple scaling relation for spheroid velocities, which captures the experimentally observed trends. These findings place magnetic spheroid assembly within the broader class of overdamped dissipative particulate systems, analogous to granular flows and dense suspensions under external forcing [23, 24]. In tissue engineering, such dissipative, stress-relaxing dynamics can support self-organization, morphogenesis, and matrix remodeling. Mechanical compaction and confinement are known to guide microtissue fusion, extracellular matrix organization, and differentiation via mechanosensitive signaling. Here, magnetic forces could add remote, programmable control over spheroid positioning, collective rearrangement, and local mechanical stimulation, complementing self-organization and bioprinting approaches [25, 26, 27]. Furthermore, the emergence of ring-like structures at high magnetic forcing demonstrates that assembloid morphology can be governed directly by the spatial structure of the magnetic force field. At small magnet-well surface separations, reversal of the tangential magnetic force above the magnet center creates an outward-directed transport region surrounded by inward attraction, generating a radial trapping zone near the magnet edge that stabilizes ring-like assembloids. As spheroid number increases, the comparatively stronger inward forces acting on outer spheroids destabilize this balance and drive collapse toward the magnet center. These results illustrate how even simple magnet geometries can generate nontrivial assembly patterns through the exploitation of spatially varying magnetic force landscapes, providing a route toward programmable design in magnetic biofabrication.

Beyond dynamics and pattern formation, our results show how collective stresses emerge during magnetic assembly and can be controlled through system geometry. Magnet shape and spheroid number determine both the magnitude and spatial distribution of compressive stresses within the assembloid. Cylindrical magnets concentrate stresses near the center, whereas alternative geometries redistribute them more uniformly. Radial and vertical stresses arise through distinct mechanisms: radial stresses emerge collectively through inter-spheroid interactions, whereas vertical stresses are primarily determined by the magnetic loading of individual spheroids. Biofabrication parameters such as spheroid number, MNP loading, and magnet design therefore provide control over interaction-dominated and load-dominated regimes. The predicted compressive stresses are on the order of 10^2^ Pa and should be interpreted in the context of early tissue formation. At this stage, spheroids remain predominantly cell-rich aggregates with limited ECM deposition and relatively low stiffness (few hundred Pa), and the assembloid itself is more appropriately viewed as a porous assembly of interacting spheroids than as a mechanically continuous tissue. These stress levels remain substantially below cyclic hydrostatic and compressive loading protocols commonly used for MSC mechanostimulation, which are typically in the kPa range [28, 29, 30], but are larger than fluid shear stresses frequently reported to regulate MSC behavior, often around 1 Pa [31, 32]. Because the stiffness of these early spheroids is itself only a few hundred Pa, stresses of this magnitude are expected to generate relevant local deformations within the aggregates [33]. Therefore the measured stresses may still be biologically relevant because MSC differentiation is known to respond to mechanical stimulation, with tensile strain favoring osteogenesis, compressive loading favoring chondrogenesis, and shear stress upregulating osteogenic genes [34, 35]. Importantly, the present simulations do not represent sustained mechanostimulation, which would require accounting for evolving material properties, active cellular forces, and progressive fusion of the assembloid [22]. They may nevertheless establish biophysically relevant initial conditions that influence subsequent fusion and maturation.

The minimal character of the model also introduces several limitations. Magnetic forces are estimated assuming a fixed fraction of nanoparticle incorporation, and spatial heterogeneity in magnetization is not explicitly resolved. Interfacial interactions are described using an effective parameter, without resolving microscopic adhesion, poroelastic effects, or deformation at the spheroid–well surface interface. In the same vein, particle rotation, including rolling, is not explicitly modeled but instead captured implicitly through an effective interfacial drag. Fluid resistance is furthermore approximated using Stokes drag, neglecting hydrodynamic corrections arising from wall proximity and neighboring spheroids, although at the large magnetic forces considered here medium drag is expected to play only a minor role in the overall dynamics. In addition, spheroids are treated as elastic spheres, neglecting large deformation, growth, and active cellular processes that may become relevant at longer timescales. Despite these simplifications, the model captures the key experimental observations, suggesting that magnetic bioassembly can be reduced to a minimal set of governing physical mechanisms. Future work could extend this framework by incorporating deformability, time-dependent material properties, and feedback between mechanical stresses and biological responses, enabling predictive control over both structure and function in engineered tissues.

## 4. Conclusion

We demonstrate that magnetic spheroid assembly can be modeled as an overdamped particulate process in which collective behavior emerges from a minimal set of physical interactions. Combining experiments and an individual-based model, we show that assembly dynamics are governed primarily by magnetic forcing and effective interfacial drag, while nontrivial aggregation patterns such as ring-like structures arise directly from the spatial pattern of the magnetic force field. Beyond structural organization, magnetic assembly generates spatially heterogeneous stress fields that are strongly controlled by magnet geometry and spheroid number, with a clear distinction between collective radial stresses and single-particle vertical loading. These findings establish a predictive physical framework for magnetic biofabrication, demonstrating how even simple magnetic geometries can be exploited to program both tissue architecture and mechanical microenvironment in engineered living systems.

## 5. Methods

### 5.1. Experimental methods

#### 5.1.1. hPDCs 2D expansion

Human periosteum-derived cells (hPDCs) were isolated from periosteal biopsies obtained from multiple donors and pooled to generate donor-independent cell populations. The hPDC pools were expanded up to passage 9 under controlled culture conditions (37 °C, 5 % CO_2_, and 95 % relative humidity) in Dulbecco’s modified Eagle medium (DMEM; Life Technologies, UK) supplemented with 10 % fetal bovine serum (HyClone FBS, Thermo Scientific, USA), 1 % antibiotic–antimycotic solution (100 U mL^−1^ penicillin, 100 µg mL^−1^ streptomycin, and 0.25 µg mL^−1^ amphotericin B), and 1 *×* 10^−3^ M sodium pyruvate (Life Technologies, UK). Cells at passage 9 were subsequently used for *in vitro* experiments. All donors provided written informed consent, and all procedures were approved by the Ethical Committee for Human Medical Research of KU Leuven (approval number ML7861).

#### 5.1.2. Generation of magnetic spheroids

Microwell culture plates (AggreWell™400 or AggreWell™800; STEMCELL Technologies Inc., Canada) were treated with anti-adherence rinsing solution (STEMCELL Technologies Inc., Canada) in accordance with the manufacturer’s protocol. Human periosteum-derived cells (hPDCs) were expanded in two-dimensional culture for seven passages, after which they were rinsed with phosphate-buffered saline (PBS), detached using TrypLE Express (Life Technologies, UK), and seeded into the microwells at a concentration of 300,000 cells well^−1^. Upon seeding, the cells spontaneously aggregated to generate three-dimensional microtissues. Microtissues were maintained under chondrogenic differentiation conditions for either 3 or 7 days using a chemically defined medium (C8). The C8 formulation consisted of low-glucose Dulbecco’s modified Eagle medium (LG-DMEM; Gibco) supplemented with 1 % antibiotic–antimycotic (100 U mL^−1^ penicillin, 100 µg mL^−1^ streptomycin, and 0.25 µg mL^−1^ amphotericin B), 1 mM ascorbate-2-phosphate, 100 nM dexamethasone, 40 µg mL^−1^ proline, and 20 µM of the Rho-associated protein kinase inhibitor Y27632 (Axon Medchem). In addition, the medium was supplemented with ITS+ Premix Universal Culture Supplement (Corning), providing 6.25 µg mL^−1^ insulin, 6.25 µg mL^−1^ transferrin, 6.25 ng mL^−1^ selenious acid, 1.25 µg mL^−1^ bovine serum albumin (BSA), and 5.35 µg mL^−1^ linoleic acid. Growth factors were added to the differentiation medium at the following concentrations: 100 ng mL^−1^ BMP-2 (INDUCTOS), 100 ng mL^−1^ GDF5 (PeproTech), 10 ng mL^−1^ TGF-*β*1 (PeproTech), 1 ng mL^−1^ BMP-6 (PeproTech), and 0.2 ng mL^−1^ FGF-2 (R&D Systems). Medium replacement was performed every second day until day 2 or day 6. At these time points, magnetic nanoparticles (MNPs), consisting of high-purity iron oxide (Fe_3_O_4_) nanoparticles (99.55% purity, Nanografi Nanotechnology), were prepared in C8 medium and added to the microtissues for overnight incubation at a final concentration of 30 µg Fe mL^−1^. Magnetic biofabrication experiments were subsequently conducted on day 3 or day 7. A schematic overview of the experimental timeline is provided in Figure 1A.

#### 5.1.3. Magnetic-guided biofabrication

After microtissue maturation up until day 3 or 7 in the microwells and their subsequent incubation with the MNPs, a magnetic plate (3D-printed) with cylindrical neodynium magnets (Supermagnete, N45) was used to guide the magnetic driven biofabrication of magnetized tissue spheroids. Spheroids were aspirated from the Aggrewell by pipetting and were transferred to another 24-well plate for suspension culture (GREINER) which was then placed on top of the magnetic plate for a duration of 60 seconds, allowing the spheroids to assemble into an assembloid, this can be seen in (Fig. 1C).

#### 5.1.4. Measurement of spheroid size

To make the computational model as accurate as possible, we collected data on the size and number of spheroids under A400 (AggreWell™400) and A800 (AggreWell™800) conditions at day 3 and day 7. The final assembloid size also depends on the number of spheroids loaded into the non-adherent well. Because we used two microwell types (A400 with 400 µm well edges and A800 with 800 µm well edges, the latter being deeper), the number of spheroids produced per condition differed despite identical initial cell seeding densities. Additionally, some spheroids escaped their microwells and fused with neighboring ones. For each condition, we took a microscopic image of an entire Aggrewell just before using the spheroids in magnetic field aggregation experiments. One well of each type (A400 and A800) was imaged for both time points. To identify the spheroids in these images, we trained an object detection algorithm using ilastik. The model was trained on a few selected regions of the well and then applied to the full image to separate spheroids from the background. For each detected spheroid, we measured the convex hull area, since spheroids can be approximated as spheres without holes. This area was then converted to an equivalent circle, and the radius of that circle was calculated so that each spheroid could be represented as a sphere with that radius.

#### 5.1.5. Mechanical characterization of spheroids

Compaction dynamics toward a single construct depend on the mechanical stiffness of the spheroids. The mechanical properties of spheroids, under A800 condition and without the addition of magnetic nanoparticles (MNPs), were evaluated on day 3 and 7 of chondrogenic differentiation using a fibre-optic-based Chiaro nanoindenter (Optics11 Life), mounted on an inverted microscope (Axio Observer, Zeiss). A spherical glass probe with a tip radius of 49.5 µm, attached to a cantilever with a stiffness of 0.53 N/m (Optics11 Life) was used for all measurements. Spheroids were immobilized on polystyrene Petri dishes (Corning) coated with poly-D-lysine (0.1 mg/mL; Gibco™, Thermo Fisher Scientific). Spheroids were pipetted onto the coated surface in culture medium and allowed to sediment and adhere for 15 min. The medium was then aspirated and replaced with room-temperature DPBS without calcium and magnesium (Gibco™, Thermo Fisher Scientific). Indentations were performed in the apical region of each spheroid, with the probe positioned as close as possible to the spheroid center to minimize variability arising from changes in contact geometry between the spherical probe and the curved spheroid surface. Multiple indentations were obtained within this region to account for the practical limitations of precise probe positioning. Adhesion mode was selected to improve the reliability of the initial contact point determination and prevent the indentation curve from starting in contact. Following automatic contact detection by the instrument software, the probe was retracted, held for 3 s, and repositioned approximately 10 µm above the sample surface before indentation. The indentations were performed in a displacement control mode, with a target indentation depth ≥ 5 µm. Per experimental condition, 8 spheroids were tested, with a minimum of 6 indentation curves obtained per spheroid. Each indentation consisted of a 2 s loading phase, followed by a 1 s holding phase, and a 2 s unloading phase. Curves were analyzed using Matlab® software (v. R2023a). To extract the effective Young’s modulus, the loading portion of the force–indentation curves was fitted using the Hertz contact model within the elastic range of the indentation response [36, 37, 38], assuming a Poisson’s ratio of 0.5 [39]. Effective Young’s modulus obtained at different locations of the same spheroid was averaged to report one representative value per spheroid.

### 5.2. Computational Model

In the particle-based model of spheroid assembly, each tissue spheroid is represented as a single spherical particle with only translational degrees of freedom. A semi-implicit numerical integration scheme was used to simulate the assembly process using overdamped equations of motion. The dynamics of spheroid *i* with position **x**_*i*_, velocity **v**_*i*_ and radius *R*_*i*_ are given by:

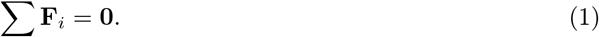

The model considers repulsive contact forces, drag forces, gravity, buoyancy, magnetic forces and stochastic forces (see Fig. 2).

#### 5.2.1. Forces

*Magnetic Field Forces*. To assign magnetic forces to magnetized spheroids, we calculate the magnetic field **H** produced by the permanent cylindrical magnet inside the well. In this case, the same approach as described by Wirthl et al. [40] was used. For a cube shaped magnet the approach of [41] was used. A 50*×*50*×*50 grid is defined over the well region. At each grid point *g*, the magnetic field is computed, allowing the magnetic force acting on a single magnetic nanoparticle to be determined at that location. For an arbitrary spheroid position within the grid, the magnetic force on a single MNP is then obtained by linear interpolation between the surrounding grid points. The total magnetic force acting on a spheroid coated with multiple MNPs is subsequently given by [40]

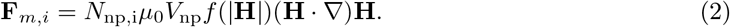

With *µ*_0_ the vacuum permeability and *V*_np_ is the volume of the nanoparticle, *N*_np,i_ is the number of nanoparticles in spheroid *i* and the magnetization model *f* (|***H***|) is defined by:

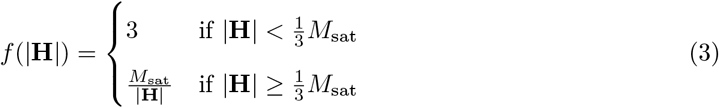

given the known particles’ saturation magnetization *M*_sat_. Finally, ***H*** denotes the magnetic field strength. The total mass of nanoparticles added (*m*_np,tot_) to all spheroids is known. Using this mass, the total number of nanoparticles

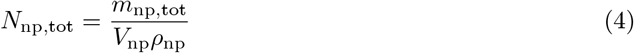

can be calculated. The number of nanoparticles on the surface of the spheroid can be found by scaling the number of nanoparticles with the area of the individual spheroid, where *N*_np,tot_ is the total number of nanoparticles and *ρ*_np_ is the density of the nanoparticles. It is assumed that not all of the nanoparticles are absorbed by the spheroids. Therefore an absorption factor *α* is defined. Thus, the number of nanoparticles on spheroid *i* is defined by:

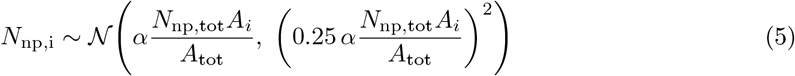

Where *A*_*i*_ represents the surface area of spheroid *i, A*_tot_ denotes the total surface area of all spheroids, and *α* is the absorption factor. The number of nanoparticles in spheroid *i*, denoted by *N*_np,i_, is modeled as a normally distributed random variable with mean proportional to its surface area and a standard deviation equal to 25% of the mean.

##### Repulsive contact force

We model repulsive contact forces between spheroids, as well as between spheroids and the bottom of the well (well surface), which is represented by a triangulated surface. For contact between spheroid *i* and spheroid *j*, the overlap distance is defined as

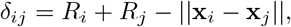

with corresponding contact normal unit vector

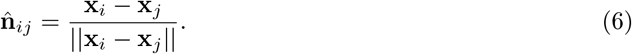

Similarly, for contact between the cylindrical well with outward normal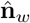 and spheroid *i*, the overlap distance is

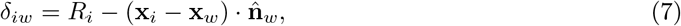

with **x**_*w*_ a reference point on the surface, and the contact normal given by 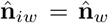. Repulsive forces are computed for all interacting pairs with overlap distance *δ* ≥ 0. The interaction force is modeled using Hertzian contact mechanics,

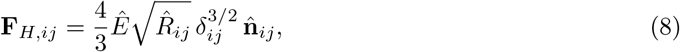

where the effective contact radius is

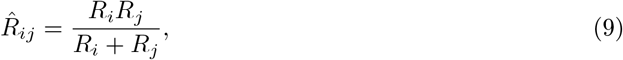

and *Ê* is the effective contact modulus, we set the effective contact modulus to the measured spheroid stiffness from NI, effectively assuming that the well surface is rigid and the spheroid poisson’s ratio approaches zero. Hertzian contact provides a minimal description of elastic repulsion between deformable spheres under compression. Although tissue spheroids are viscoelastic and may exhibit weak adhesive interactions, the magnetic loading in our system places contacts in a strongly compressed regime, where adhesive corrections to the contact area are expected to be small, and the response approaches the Hertzian limit [42]. The contact area *S*_*ij*_ used in the aggregation model is then obtained from the Hertzian contact radius, yielding

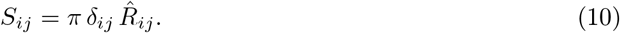

##### Interfacial friction with the well surface

Interfacial resistance between spheroids and the well surface is modeled using an effective viscous traction law that represents dissipation associated with relative motion at the contact interface. The tangential force exerted by the well surface on spheroid *i* is given by

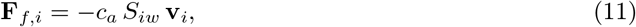

where *S*_*iw*_ denotes the spheroid-well surface contact area, **v**_*i*_ is the velocity of spheroid *i* relative to the well surface, and *c*_*a*_ is an effective tangential interfacial drag coefficient with units of [Pa s mm^−1^]. This formulation assumes that tangential resistance arises from viscous dissipation within the contact region and therefore scales linearly with velocity. The dependence on *S*_*iw*_ reflects the distributed nature of the interfacial dissipation, such that an increase in the contact area leads to a proportional increase in tangential resistance. The contact area *S*_*iw*_ is estimated using Hertzian contact mechanics and depends on the normal magnetic load acting on the spheroid. As a consequence, the magnitude of the tangential resistance exhibits a sublinear dependence on the normal force through the contact-area scaling. This sublinear scaling is consistent with single-asperity contact arguments for non-adhesive interfaces, where friction arises from distributed microscopic contacts and does not increase linearly with normal load [43].

##### Medium drag force

The drag force exerted by the surrounding medium on a spheroid is described by Stokes’ law,

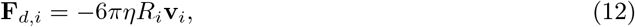

where *η* denotes the viscosity of the medium. In our system, this hydrodynamic drag is significantly smaller than the effective interfacial resistance associated with spheroid–well surface interactions and therefore does not govern lateral motion once spheroids are in contact with the well surface. However, it plays an important role during the initial phase of the experiment, where it controls sedimentation and the rapid magnetic acceleration of spheroids as they descend through the medium toward the well surface.

##### Gravitational Force

The combined gravitational and buoyancy force is

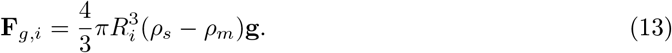

with *ρ*_*s*_ the density of the spheroids and *ρ*_*m*_ the density of the medium.

#### 5.2.2 Equations of Motion

Combining all these forces together into one equation of motion for particle *i* which is given by:

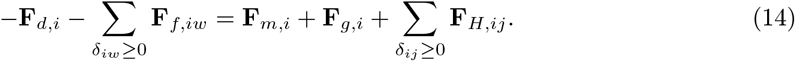

The left hand side of the equation comprises all velocity dependent forces, while the right hand side contains the body and contact forces. The equation of motion for all particles can then be written as a system of linear equations

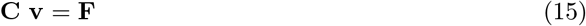

since all terms depending on velocity are linear. Here **C** is a 3n *×* 3n matrix where the diagonal elements are formed by the medium drag and the other elements are formed by the contact pairs *i* and *j* and **v** = [**v**_1_, …, **v**_*i*_, …, **v**_*n*_]^*T*^ and **F** = [**F**_1_, …, **F**_*i*_, …, **F**_*n*_]^*T*^ are the state vectors. **C** is a sparse, positive definite and a symmetric matrix, therefore this system can be solved by the conjugate gradient method.

#### 5.2.3. Time Integration

The linear system with the equations of motion is solved using semi-implicit time integration. The particle velocities are updated by solving:

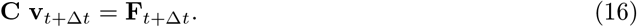

for velocity and then updating positions as

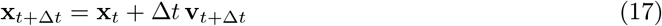

#### 5.2.4. Mechanical Stress

The mechanical stress experienced by each spheroid at its position within the simulated assembloid is determined using the virial stress theorem [44, 45]. For spheroid *i*, the local average stress tensor ***σ***_*i*_ is evaluated from inter-spheroid interaction forces and the spheroid volume. The per-spheroid virial stress is calculated as

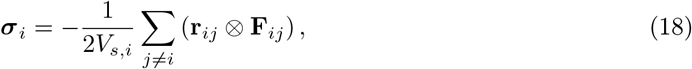

where *V*_*s,i*_ is the spheroid volume, **r**_*ij*_ = **x**_*j*_ − **x**_*i*_ is the relative position vector, and **F**_*ij*_ is the force exerted on spheroid *i* by spheroid *j* or the well surface. The factor 1*/*2 avoids double counting of pairwise interactions, and the negative sign ensures consistency with the Cauchy stress convention. The resulting tensor is decomposed into radial and vertical components. The radial mechanical stress is defined as

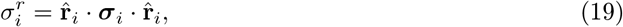

where the radial unit vector is computed by projecting the position vector onto the xy-plane,

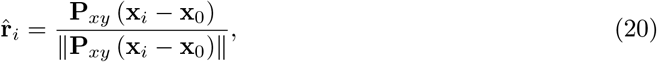

with **x**_0_ denoting the center of the magnetic field and **P**_*xy*_ the projection operator onto the xy-plane. In addition to the radial component, the vertical stress is defined as the projection of the stress tensor along the *z*-direction,

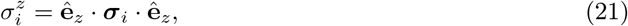

where **ê**_*z*_ is the unit vector in the vertical (*z*) direction. This component captures the mechanical stress acting perpendicular to the *xy*-plane. Mechanical stresses are evaluated for each spheroid individually and subsequently averaged over radial bins based on their distance from the center, yielding a spatially resolved stress profile across the assembloid.

### 5.3. Simulation setup

In the particle-based model of spheroid assembly, each tissue spheroid is represented as a single spherical particle. The simulation takes place inside a rigid container that mimics the 3D geometry of a single well from a 24-well suspension culture plate (GREINER). Spheroid sizes and spheroid counts are taken from Fig.1E. From these values, the average spheroid radius per condition is calculated. Assuming a standard deviation of 0.15 times the average spheroid size, spheroid sizes are then sampled from a normal distribution. Spheroids are first randomly placed within the well and allowed to sediment under gravity for 20 seconds. After this sedimentation phase, a magnetic field is applied to drive their assembly. The magnetic force acting on each spheroid is derived from force calculations performed for a single magnetic nanoparticle (MNP) on a predefined 3D grid covering the entire well volume. The particles used are superaparamagnetic ironoxide magnetic nanoparticles with radius of 22.5 nm. The magnetic force calculations assume a cylindrical neodymium magnet (N45 grade), magnetized along the negative z-axis, positioned directly beneath the well. The magnet used has a diameter of 4.76 mm and a length of 4.7 mm. At any spheroid’s position, the magnetic field intensity is obtained by linearly interpolating between the nearest grid points. The resulting force is then multiplied by the number of MNPs internalized by that spheroid to obtain the total magnetic force.

**Table 1.** Model, material, magnetic, and simulation parameters used in this study, unless otherwise specified.

| Parameter | Symbol | Value | Units | Source / rationale |
| --- | --- | --- | --- | --- |
| **General constants** |  |  |  |  |
| Magnetic permeability of vacuum | $\mu_0$ | 1.2566 | $\mu\text{N A}^{-2}$ | Fundamental constant |
| Medium viscosity | $\eta$ | 1.002 | mPa s | Approximated as water at room temperature |
| Spheroid density | $\rho_s$ | 1014 | $\text{kg m}^{-3}$ | Reported value for cellular aggregates [22] |
| Medium density | $\rho_m$ | 1000 | $\text{kg m}^{-3}$ | Density of water |
| Viscous well surface drag constant | $c_a$ | 33.2363 | $\text{Pa s mm}^{-1}$ | Derived from experiments Fig. 2D |
| Effective contact modulus | $\hat{E}$ | [254, 361] | Pa | Experimentally measured Fig. 1D–E |
| Number of spheroids | $N_s$ | [1159, 409, 294, 261] | – | Measured from experimental conditions Fig. 1F |
| Average spheroid radius | $\bar{R}$ | [92, 120, 139, 160] | $\mu\text{m}$ | Experimentally measured 1F |
| Spheroid size standard deviation | $\sigma(R)/\bar{R}$ | 0.15 | – | Assumed value |
| **Magnet properties** |  |  |  |  |
| Magnet separation | $d_{\text{mag}}$ | 0.42 | mm | Fixed experimental geometry |
| Magnet radius | $R_{\text{mag}}$ | 2.38 | mm | Manufacturer specifications |
| Magnet length | $L_{\text{mag}}$ | 4.76 | mm | Manufacturer specifications |
| Magnet magnetization | $M$ | $1.07 \times 10^9$ | $\text{A mm}^{-1}$ | Manufacturer specifications |
| **Nanoparticle properties** |  |  |  |  |
| Nanoparticle radius | $R_{\text{np}}$ | 10.75 | nm | Manufacturer specifications |
| Nanoparticle density | $\rho_{\text{np}}$ | 5.24 | $\text{g cm}^{-3}$ | Manufacturer specifications |
| MNP absorption factor | $\alpha$ | 0.56 | – | Literature value [22] |
| MNP saturation magnetization | $M_{\text{sat}}$ | $3.11 \times 10^8$ | $\text{A mm}^{-1}$ | Literature value [22] |
| Total MNP mass | $m_{\text{np,tot}}$ | [10, 20] | $\mu\text{g}$ | Experimental loading conditions |
| **Simulation parameters** |  |  |  |  |
| Simulation timestep | $\Delta t$ | 5 | $\mu\text{s}$ | Chosen for numerical stability |
| Settling timestep | $\Delta t_{\text{set}}$ | 5 | ms | Chosen for efficient equilibration |
| Settling time | $t_{\text{set}}$ | 40 | s | Selected to match experiment |
| Simulation time | $t_{\text{sim}}$ | 80 | s | Covers full experimental timescale |
| Grid resolution | – | $50 \times 50$ | – | Compromise between accuracy and cost |
| Well diameter | $d_{\text{well}}$ | 16 | mm | Experimental geometry |
| Well height | $h_{\text{well}}$ | 15 | mm | Experimental geometry |

## Acknowledgments

This research was supported by the Research Foundation Flanders (FWO) under project G042425N. acknowledges support from FWO through fellowship 1SD7225N, and D.L. acknowledges support from FWO through fellowship 1SA4426N. Additional support was provided by the KU Leuven Internal Funds (project IDs ID-N/3M230283 and C24M/22/058, I.P.). This work was carried out within Prometheus, the KU Leuven R&D division for skeletal tissue engineering (www.kuleuven.be/prometheus). The authors thank the staff of FIBEr, KU Leuven Core Facility for Biomechanical Experimentation (www.biomech.be/fiber), for their assistance with the nanoindentation experiments.

## Code availability

Simulations were performed using the particle-based simulation framework Mpacts. The source code related to the simulations presented in this study is available on GitLab [46].

## Data Availability

Data supporting the findings of this study are available from the corresponding author upon reasonable request.

## Author contributions

M.G.: Conceptualization, Methodology, Software, Validation, Formal analysis, Investigation, Writing - Original Draft K.I.: Conceptualization, Methodology, Investigation K.Z.D.: Formal analysis, Investigation, Writing - Original Draft M.T.: Formal analysis, Investigation, Writing - Original Draft D.L.: Resources, Writing - Original Draft G.S.: Writing - Review & Editing, Supervision D.S.: Writing - Review & Editing, Supervision, Funding acquisition I.P.: Conceptualization, Methodology, Writing - Review & Editing, Supervision, Funding acquisition B.S.: Conceptualization, Methodology, Software, Formal analysis, Writing - Original Draft, Supervision, Funding acquisition

## Appendix A. Estimation of sliding friction from experiments

We consider a force balance that combines area-dependent sliding friction with Stokes drag. Close to the magnet, the normal force is large, leading to an increased contact area and thus dominant sliding friction. Far from the magnet, magnetic forces are weaker and Stokes drag cannot be neglected. Under these assumptions, the tangential force balance reads:

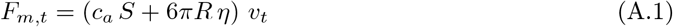

The spheroid–well contact area *S* is estimated using a Hertzian contact model for a sphere pressed against a rigid flat surface. The contact area is given by *S* = *πRδ*, where the indentation depth *δ* follows:

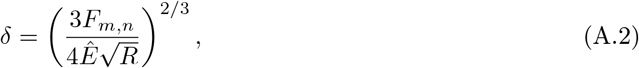

with *Ê* the effective elastic modulus. Substituting this expression into Eq. (A.1) yields a model for the tangential velocity:

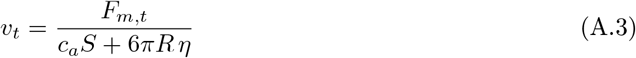

or, equivalently,

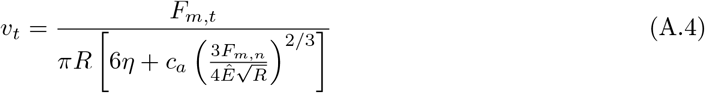

We estimate the sliding friction coefficient *c*_*a*_ by fitting this expression to experimentally measured tangential velocities *v*_*t*_, using characterized or independently estimated values of *F*_*m,t*_, *F*_*m,n*_, *R, Ê*, and *η*. For each tracked spheroid, the measured spheroid radius *R* is used directly in the force balance and contact-area estimation.

## Appendix B. Scaling of sliding velocity with spheroid size and MNP loading

We consider a fixed total volume of spheroids *V*_*s*,tot_ (defined by the total cell mass used in the experiment), composed of identical spheroids of radius *R* and volume 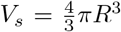. The number of spheroids is then *N*_*s*_ = *V*_*s*,tot_*/V*_*s*_. A fixed fraction *α* of the total number of magnetic nanoparticles *N*_np,tot_ is distributed uniformly across all spheroids. Since the nanoparticle size and density are fixed, *N*_np,tot_ ∝ *m*_np,tot_. Thus the number of nanoparticles per spheroid is

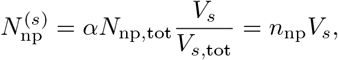

with spheroid-incorporated magnetic particle density *n*_np_ = *αN*_np,tot_*/V*_*s*,tot_. The total magnetic force on a spheroid is then:

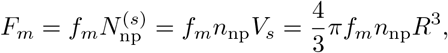

from which it follows that *F*_*m*_ ∝ *n*_np_*R*^3^. We decompose this force into a normal and tangential component,

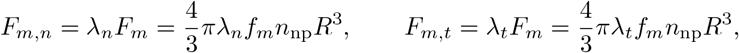

with 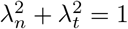. For a Hertz-like contact force, the normal force balance reads

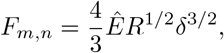

so that

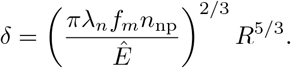

Thus, 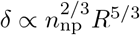. The Hertz contact area follows as

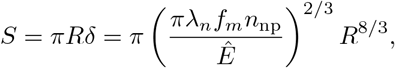

and therefore 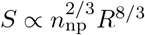. Let the sliding resistance contain both Stokes drag in the surrounding liquid and an area-dependent friction with coefficient *c*_*a*_. The tangential force balance is then

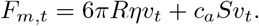

Substitution gives

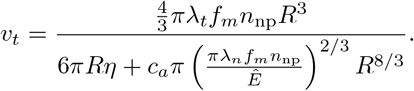

This may be written as the scaling relation

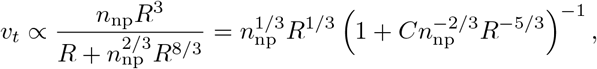

where *C* collects constants independent of *n*_np_ and *R*. In the low-loading limit, where Stokes drag dominates, this gives

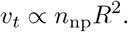

In the high-loading limit, where area-dependent friction dominates, the correction term vanishes and we recover

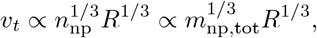

where the latter relation follows from the assumption of fixed total spheroid volume *V*_*s*,tot_.

