## Supplementary Information for "Modeling spheroid assembly dynamics and mechanical stress generation in magnetic-based biofabrication"

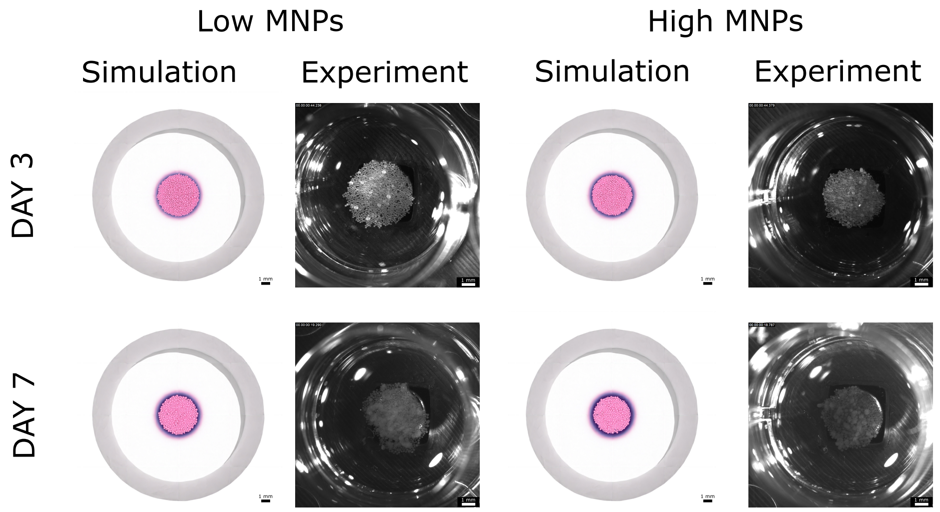


**Supplementary Fig. S1:** Representative images of the final assembloid morphology after 60 seconds in both simulation and experimental conditions. Comparisons are shown for low and high MNP loading, as well as for low MNP loading in Day 3 and Day 7 spheroids. All assembloids were generated using spheroids formed in an A400 well plate.

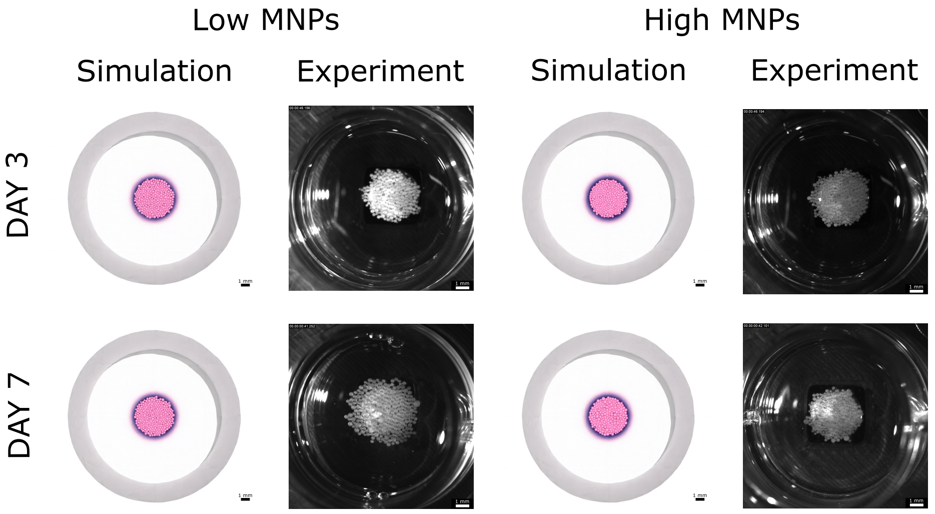


**Supplementary Fig. S2:** Representative images of the final assembloid morphology after 60 seconds in both simulation and experimental conditions. Comparisons are shown for low and high MNP loading, as well as for low MNP loading in Day 3 and Day 7 spheroids. All assembloids were generated using spheroids formed in an A800 well plate.

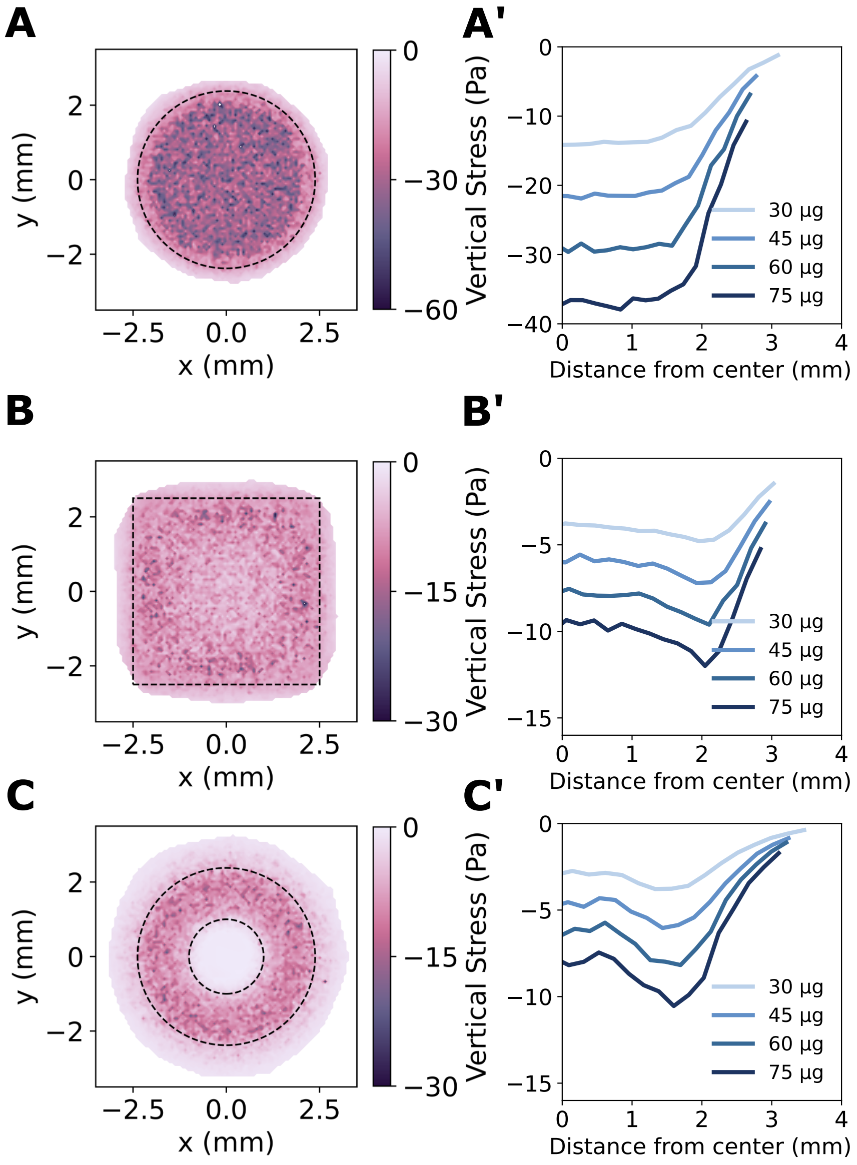


**Supplementary Fig. S3:** (A–C) Magnetic field maps generated by cylindrical **(A)**, ring-shaped **(B)**, and square **(C)** magnets used to drive spheroid assembly, averaged over 50 independent repeats. (A'-C') Spatial distributions of the coarse-grained vertical stress within the assembled spheroid aggregates, averaged over 50 independent repeats. (A"-C") Vertical stress profiles as a function of distance from the center for increasing MNP content (30–75 µg), averaged over 50 independent repeats.

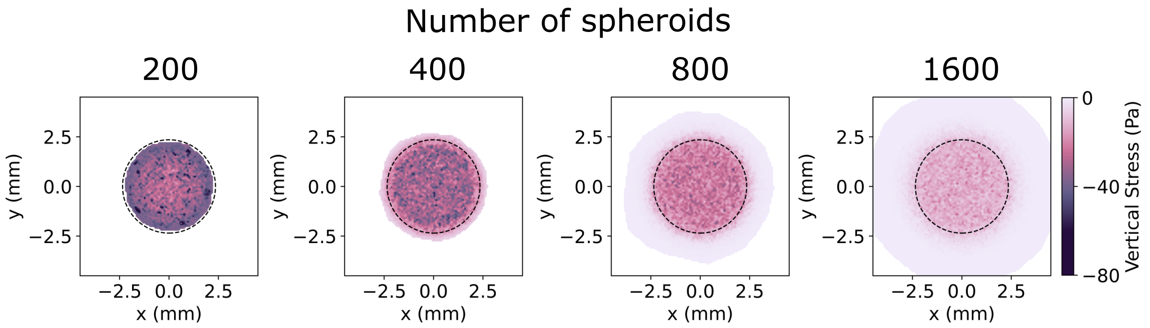


**Supplementary Fig. S4:** Radial stress maps for assemblies composed of 200, 400, 800, and 1600 spheroids, respectively, with total MNP content held constant, averaged over 50 independent repeats.
